# Cytoprotective effects of (E)-N-(2-(3, 5-dimethoxystyryl) phenyl) furan-2-carboxamide (BK3C231) against 4-nitroquinoline 1-oxide-induced damage in CCD-18Co human colon fibroblast cells

**DOI:** 10.1101/777193

**Authors:** Huan Huan Tan, Noel F. Thomas, Salmaan H. Inayat-Hussain, Kok Meng Chan

## Abstract

Stilbenes are a group of chemicals characterized with the presence of 1,2-diphenylethylene. Previously, our group has demonstrated that synthesized (E)-N-(2-(3, 5-dimethoxystyryl) phenyl) furan-2-carboxamide (BK3C231) possesses potential chemopreventive activity specifically inducing NAD(P)H:quinone oxidoreductase 1 (NQO1) protein expression and activity. In this study, the cytoprotective effects of BK3C231 on cellular DNA and mitochondria were investigated in normal human colon fibroblast, CCD-18Co cells. The cells were pretreated with BK3C231 prior to exposure to the carcinogen 4-nitroquinoline 1-oxide (4NQO). BK3C231 was able to inhibit 4NQO-induced cytotoxicity. Cells treated with 4NQO alone caused high level of DNA and mitochondrial damages. However, pretreatment with BK3C231 protected against these damages by reducing DNA strand breaks and micronucleus formation as well as decreasing losses of mitochondrial membrane potential (ΔΨm) and cardiolipin. Interestingly, our study has demonstrated that nitrosative stress instead of oxidative stress was involved in 4NQO-induced DNA and mitochondrial damages. Inhibition of 4NQO-induced nitrosative stress by BK3C231 was observed through a decrease in nitric oxide (NO) level and an increase in glutathione (GSH) level. These new findings elucidate the chemopreventive potential of BK3C231 specifically its cytoprotective effects in human colon fibroblast CCD-18Co cell model.

## 1. Introduction

Cancer-related mortality has tremendously increased and is expected to further increase despite emerging medical improvements [1]. The global cancer burden is estimated to have risen in 2018 with colorectal cancer being the third most commonly diagnosed cancer and is ranked second in terms of mortality due to poor prognosis worldwide [2]. In Malaysia, cancer is the third most common cause of death after cardiovascular diseases and respiratory diseases. According to Malaysia National Cancer Registry (MNCR) Report 2007-2011, colorectal cancer is the second most common cancer among Malaysian residents [3].

Advances in costly surgical and medical therapies for primary and metastatic colorectal cancer have had limited impact on cure rates and long-term survival [4, 5]. Local recurrences after surgery, treatment-induced long-term complications and toxicities, chemotherapy-induced adverse effects as well as cancerous cell resistance towards chemotherapy due to development of multidrug resistance phenotypes and tumour heterogeneity become major reasons for chemoprevention to gain momentum [6–8].

Chemopreventive approaches have effectively decreased cancer incidence rates such as for lung cancer and cervical cancer [2], thus strengthening the need to prevent or limit the disease to occur in the first place. Cancer chemoprevention is an essential approach which uses nontoxic natural or synthetic pharmacological agents to prevent, block or reverse the multistep processes of carcinogenesis [9, 10]. Chemopreventive agents inhibit the invasive development of cancer by affecting the three defined stages of carcinogenesis namely initiation, promotion and progression which are induced by carcinogens through genetic and epigenetic mechanisms [11, 12].

Exposure of cells to carcinogen causes DNA mutation and leads to accumulation of additional genetic changes through sustained cell proliferation. This rapid and irreversible process is known as tumour initiation, the first stage of carcinogenesis. Tumour promotion, which is referred to as the lengthy and reversible second stage of carcinogenesis, involves the selective clonal expansion of initiated cells to produce preneoplastic lesion which allows for additional mutations to accumulate. The final stage of carcinogenesis, tumour progression, involves neoplastic transformation after accumulating chromosomal aberrations and karyotypic instability resulting in metastatic malignancy [13, 14].

Altered cellular redox status and disrupted oxidative homeostasis play a key role towards cancer development by enhancing DNA damage and modifying key cellular processes such as cell proliferation and apoptosis [15]. Oxidative/nitrosative stress is the result of disequilibrium between reactive oxygen species (ROS)/reactive nitrogen species (RNS) and antioxidants [16]. If oxidative/nitrosative stress persists, this may lead to modification of signal transduction and gene expression, which in turn lead to mutation, transformation and progression of cancer [17, 18].

Stilbenes are a group of phenylpropanoids produced in the skin, seeds, leaves and sapwood of a wide variety of plant species including dicotyledon angiosperms such as grapevine (*Vitis vinifera*), peanut (*Arachis hypogaea*) and Japanese knotweed (Fallopia Japonica); monocotyledons like sorghum (*Sorghum bicolor*) and gymnosperms such as several *Pinus* and *Picea* species [19–21]. They are a well-known class of naturally occurring phytochemicals acting as antifungal phytoalexins, providing protection against UV light exposure and also involved in bacterial root nodulation and coloration [19,22–24]. These compounds bear the core structure of 1,2-diphenylethylene in which two benzene rings are separated by an ethanyl or ethenyl bridge [25].

Despite being known as plant defense compounds, stilbenes have an enormous diversity of effects on biological and cellular processes applicable to human health, particularly in chemoprevention. Resveratrol, as the biosynthetic precursor of most oligostilbenoids, has been known to possess a myriad of biological activities such as anticancer, antioxidant, anti-aging, antimicrobial, cardioprotection, anti-diabetes, anti-obesity, and anti-inflammation [26–33]. However, low water solubility and poor bioavailability are the major setbacks to the exploitation of these biological activities [34, 35].

Our group has previously demonstrated that synthetic stilbene BK3C231 (Fig 1) potently induced antioxidant gene NQO1 as a detoxifying mechanism in human embryonic hepatocytes, WRL-68 cells [36]. Therefore in this study, we proposed to elucidate the chemopreventive effects specifically the cytoprotective effects of BK3C231 using normal human colon fibroblast CCD-18Co cells. We anticipate this study will accelerate the development of BK3C231 as a potential drug for chemoprevention.

**Fig 1.**
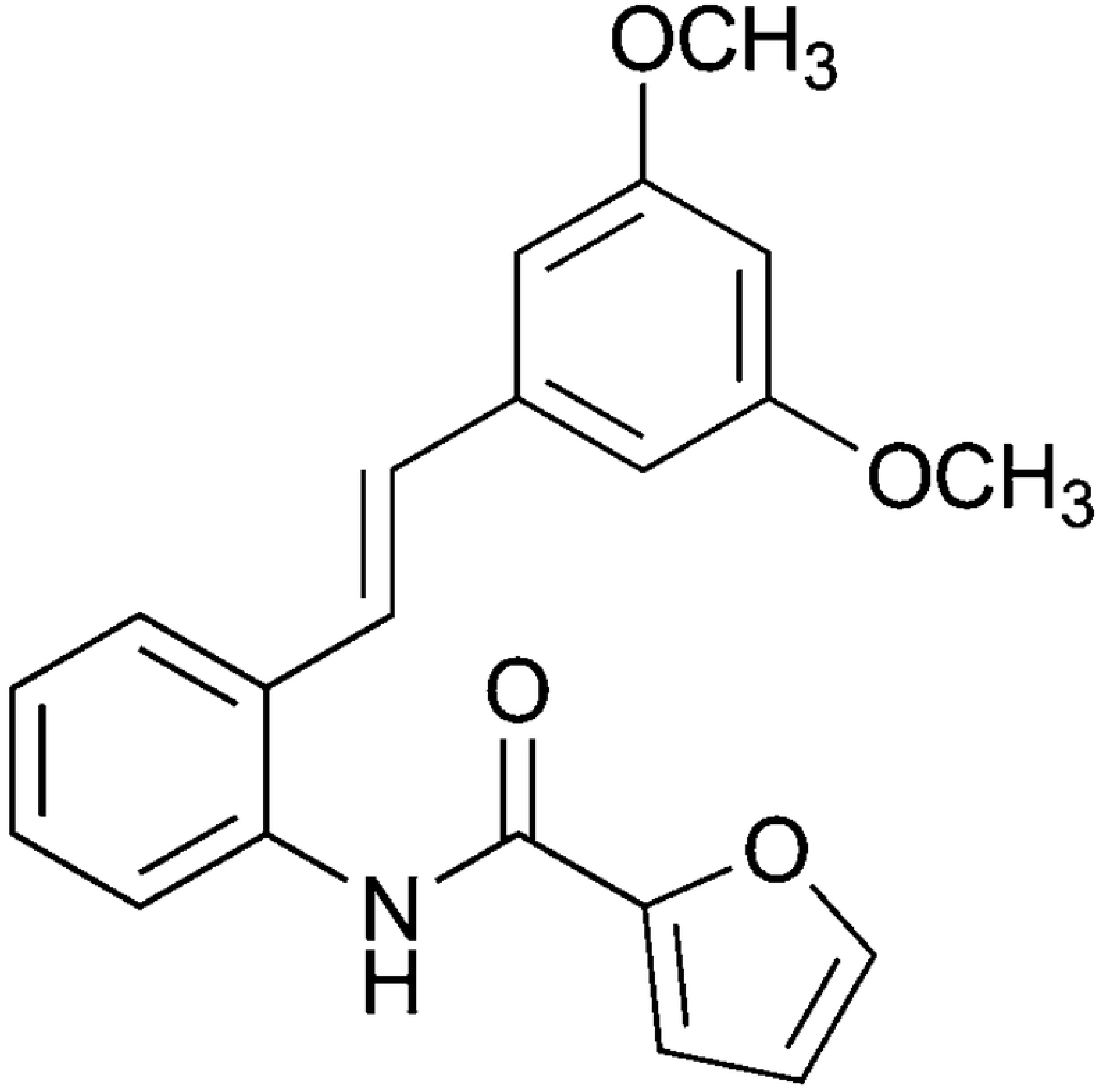
Chemical structure of BK3C231 [36].

## 2. Materials and methods

### 2.1 Test compounds

(E)-N-(2-(3, 5-Dimethoxystyryl) phenyl) furan-2-carboxamide (BK3C231) was synthesized and contributed by Dr. Noel Francis Thomas and Dr. Kee Chin Hui from Department of Chemistry, Faculty of Science, University of Malaya (Kuala Lumpur, Malaysia). 4-Nitroquinoline 1-oxide (4NQO) (Cas. No: 56-57-5, ≥98% purity) was purchased from Sigma-Aldrich (St. Louis, MO, USA). Stock solution of BK3C231 at 100mM and 4NQO at 25mg/mL were prepared by dissolving the compounds in solvent dimethyl sulfoxide (DMSO, Thermo Fisher Scientific, Waltham, MA, USA).

### 2.2 Cell culture

The normal human colon fibroblast CCD-18Co cell line (ATCC CRL-1459) was obtained from the American Type Culture Collection (ATCC, Manassas, VA, USA). CCD-18Co cells were grown in Minimum Essential Medium (MEM, Gibco, Grand Island, NY, USA) supplemented with 10% fetal bovine serum (FBS, Biowest, Nuaillé, France) and 1% 100x Antibiotic-Antimycotic solution (Nacalai Tesque, Kyoto, Japan). All cells were between passages 3–5 for all experiments and maintained at 37°C with 5% CO_2_.

### 2.3 MTT cytotoxicity assay

CCD-18Co cells were seeded in 96-well microplate (Nest Biotechnology, Jiangsu, China) at the concentration of 5 x 10^4^ cells/mL in a volume of 200 μL per well. The seeded cells were incubated under 5% CO_2_ at 37°C for 24 hours prior to respective compound treatments at different timepoints. After incubation, 20 µL of MTT (Sigma-Aldrich, St. Louis, MO, USA) solution (5mg/mL in PBS) was added to the treated cells and further incubated for 4 hours at 37°C. Subsequently, the total medium in each well was discarded and the crystalline formazan was solubilised using 200 µL DMSO. For complete dissolution, the plate was incubated for 15 minutes followed with gentle shaking for 5 minutes. The cytotoxicity of BK3C231 and 4NQO was assessed by measuring the absorbance of each well at 570 nm. Mean absorbance for each compound concentration was expressed as a percentage of vehicle control absorbance and plotted versus compound concentration. Inhibitory concentration that kills 50% of cell population (IC_50_) represents the compound concentration that reduced the mean absorbance at 570 nm to 50% of those in the vehicle control wells.

### 2.4 Alkaline comet assay

Seeded cells (5 x 10^4^ cells/mL) in 6-well plate (Nest Biotechnology, Jiangsu, China) were pretreated with BK3C231 at 6.25 μM, 12.5 μM, 25 μM and 50 μM for 2 hours prior to 4NQO treatment at 1 μM for 1 hour. Following incubation, detached cells in the medium were collected and added back to trypsinised cells. Then, the suspension was transferred to tube for centrifugation (450 x g/5 minutes at 4^◦^C). The supernatant was removed and pellet was washed with Ca^2+^- and Mg^2+^-free PBS and re-centrifuged. The pellets left at the bottom were mixed thoroughly with 80 μl of 0.6% w/v LMA (Sigma-Aldrich, St. Louis, MO, USA). The mixture was then pipetted onto the hardened 0.6% w/v NMA (Sigma-Aldrich, St. Louis, MO, USA) as the first layer gel on the slide. Cover slips were placed to spread the mixture and slides were left on ice for LMA to solidify. Following removal of the cover slips, the embedded cells were lysed in a lysis buffer containing 2.5M NaCl (Merck Milipore, Burlington, MA, USA), 1 mM Na_2_EDTA (Sigma-Aldrich, St. Louis, MO, USA), 10 mM Tris (Bio-Rad, Hercules, CA, USA) and 1% Triton X-100 (Sigma-Aldrich, St. Louis, MO, USA) overnight at 4^°^C. After lysis, the slides were soaked in electrophoresis buffer solution for 20 minutes for DNA unwinding before electrophoresis at 300 mA, 25V for 20 minutes. Subsequently, the slides were rinsed with neutralising buffer for 5 minutes and stained with 30 μL of 50 μg/mL ethidium bromide (EtBr, Sigma-Aldrich, St. Louis, MO, USA) solution. Slides were left overnight at 4^◦^C before analyzing with Olympus BX51 fluorescence microscope (Tokyo, Japan) equipped with 590 nm filter. DNA damage scoring was performed on 50 cells per slide whereby tail moment representing the product of tail length and fraction of total DNA in tail was quantified using Comet Score^TM^ software (TriTek Corp, Sumerduck, VA, USA).

### 2.5 Cytokinesis-block micronucleus (CBMN) assay

Seeded cells (5 x 10^4^ cells/mL) in 6-well plate were pretreated with BK3C231 at 6.25 μM, 12.5 μM, 25 μM and 50 μM for 2 hours prior to 4NQO treatment at 1 μM for 2 hours. After incubation, cells were treated with 4.5 μg/mL Cytochalasin B (Sigma-Aldrich, St. Louis, MO, USA) for 24 hours to block cytokinesis. The cells were then harvested and centrifuged (450 x g/5 minutes at 4^◦^C). The supernatant was removed and pellet was resuspended with 300 μL of 0.075M KCl solution for 5 minutes. The cells were then fixed with Carnoy’s solution consisting of acetic acid (Sigma-Aldrich, St. Louis, MO, USA) and methanol (HmbG Chemicals, Hamburg, Germany) prepared at the ratio of 1:3 and spreaded on glass slides which were placed on a slide warmer. The slides were dried overnight and stained with 30 μL of 20 μg/mL acridine orange (AO, Sigma-Aldrich, St. Louis, MO, USA) prior to fluorescence microscopic observation. The number of viable mononucleated, binucleated and multinucleated cells per 500 cells were scored to derive Nuclear Division Index (NDI) and frequency of micronucleus in 1,000 binucleated cells was measured.

### 2.6 Mitochondrial membrane potential (ΔΨm), mitochondrial mass and ROS assessment

The treated cells (5 x 10^4^ cells/mL) were collected by centrifugation (450 x g/5 minutes at 4^◦^C). The supernatant was discarded and pellet was resuspended with 1 mL fresh prewarmed FBS-free MEM with addition of 1 μL of 50 μM tetramethylrhodamine ethyl ester (TMRE, Thermo Fisher Scientific, Waltham, MA, USA), 5 mM nonyl acridine orange (NAO, Sigma-Aldrich, St. Louis, MO, USA), 10 mM hydroethidine (HE, Thermo Fisher Scientific, Waltham, MA, USA) or 10 mM 2’,7’-dichlorodihydrofluorescein diacetate (DCFH-DA, Thermo Fisher Scientific, Waltham, MA, USA). The cells stained with TMRE or NAO were incubated for 15 minutes at 37^◦^C whereas cells stained with HE and DCFH-DA were incubated for 30 minutes at 37^◦^C in the dark. After incubation, the cells were centrifuged (450 x g/5 minutes at 4^◦^C) and pellet was washed with 1 mL chilled PBS solution. The supernatant was discarded and 500 μL of chilled PBS was used to resuspend the pellets. The stained cell suspension was transferred to flow tubes and analyzed using FACSCanto II Flow Cytometer (BD Biosciences, San Jose, CA, USA).

### 2.7 Intracellular nitric oxide (NO) assessment using BD Pharmingen™ Orange Nitric Oxide (NO) Probe staining

The seeded cells (5 x 10^4^ cells/mL) were pre-stained with 1 μL Orange NO probe (BD Biosciences, San Jose, CA, USA) per 500 μL cell suspension for 30 minutes. The cells were then pretreated with BK3C231 at 50 μM for 2 hours, 4 hours, 6 hours, 12 hours and 24 hours prior to 4NQO treatment at 1 μM for 1 hour. The stained and treated cells were centrifuged (450 x g/5 minutes at 4^◦^C) and pellet was washed with 1 mL chilled PBS solution. The supernatant was discarded and 500 μL of chilled PBS was used to resuspend the pellets. The stained cell suspension was transferred to flow tubes and analyzed using FACSCanto II Flow Cytometer (BD Biosciences, San Jose, CA, USA).

### 2.8 Extracellular nitric oxide (NO) assessment using Griess reagent

CCD-18Co cells were seeded in culture dish (60 x 15 mm) at the concentration of 5 x 10^4^ cells/mL. The seeded cells were incubated under 5% CO_2_ at 37°C for 24 hours. The cells were then pretreated with BK3C231 at 50 μM for 2 hours, 4 hours, 6 hours, 12 hours and 24 hours prior to 4NQO treatment at 1 μM for 1 hour. Subsequently, 100 μL of culture medium from each sample was collected and mixed with the same volume of Griess reagent (1% sulfanilamide in 5% phosphoric acid and 0.1% N-(1-naphthyl)ethylenediamine (NNED) hydrochloride in distilled water, Merck Milipore, Burlington, MA, USA) in 96-well microplate. Absorbance of the mixture in each well was determined at 570 nm. The concentration of nitrite accumulated in the culture was determined in comparison to the sodium nitrite standards.

### 2.9 Glutathione (GSH) assessment using Ellman’s reagent

The treated cells (5 x 10^4^ cells/mL) were detached, collected and centrifuged (450 x g/5 minutes at 4^◦^C). The supernatant was discarded and pellet was resuspended in 100 μL ice-cold lysis buffer (50 mM K_2_HPO_4_, 1 mM EDTA, pH 6.5 and 0.1 % v/v Triton X-100, Sigma-Aldrich, St. Louis, MO, USA). The cells were incubated on ice for 15 minutes with gentle tapping from time to time. The crude lysates were cleared by centrifugation (10000 x g/15 minutes at 4°C). At this point, the lysates were used immediately or stored at −80°C for a day or two. Then, 50 μl of lysates and GSH standards (two fold dilution from 1.25 mM to 0 mM dissolved in reaction buffer consisting of 0.1 M Na_2_HPO_4_.7H_2_O and 1 mM EDTA, pH 6.5, Sigma-Aldrich, St. Louis, MO, USA) were pipetted into designated wells in a 96-well microplate. After adding 40 μl of reaction buffer (0.1 M Na_2_HPO_4_.7H_2_O and 1 mM EDTA, pH 8), 10 μl of 4 mg/ml 5,5′-dithiobis(2-nitrobenzoic acid) (DTNB, Sigma-Aldrich, St. Louis, MO, USA) in reaction buffer pH 8 was added to wells containing samples and standards. The plate was incubated for 15 minutes at 37°C. Absorbance of each well was measured at 405 nm using microplate reader (Bio-Rad, Hercules, CA, USA). The concentration of free thiols in samples was calculated based on GSH standard and expressed as nmol/mg protein after protein concentration was quantified using Bradford’s method.

### 2.10 Statistical analysis

The data are expressed as the mean ± standard error of mean (S.E.M.) from at least three independent experiments. The statistical significance was evaluated using one-way ANOVA with the Tukey post hoc test used to assess the significance of differences between multiple treatment groups. Differences were considered statistically significant with a probability level of p<0.05.

## 3. Results

### 3.1 Cytotoxic assessment of BK3C231 and 4NQO

The non-cytotoxic concentrations of BK3C231 and 4NQO were determined using MTT cytotoxicity assay. BK3C231 did not show evidence of cytotoxicity up to 50 µM treatment, however an IC_50_ value of 99 µM was observed (Fig 2A). Therefore a series of BK3C231 concentrations ranging from 6.25 µM till 50 µM was used for subsequent experiments. On the other hand, 4NQO treatment exerted no cytotoxicity at 1 hour. However, reduction in cell viability was significant with IC_50_ values observed starting from 2 hours till 24 hours (Fig 2B). Hence, 4NQO concentration at 1 µM was selected to induce genotoxicity and mitochondrial toxicity in subsequent experiments as used by previous studies as well [37, 38]. Interestingly, in comparison to 4NQO-treated cells whereby cell viability greatly reduced especially at higher concentrations, BK3C231 was able to suppress 4NQO-induced cytotoxicity by increasing cell viability up to 8-fold with no IC_50_ value observed (Fig 2C).

**Fig 2.**
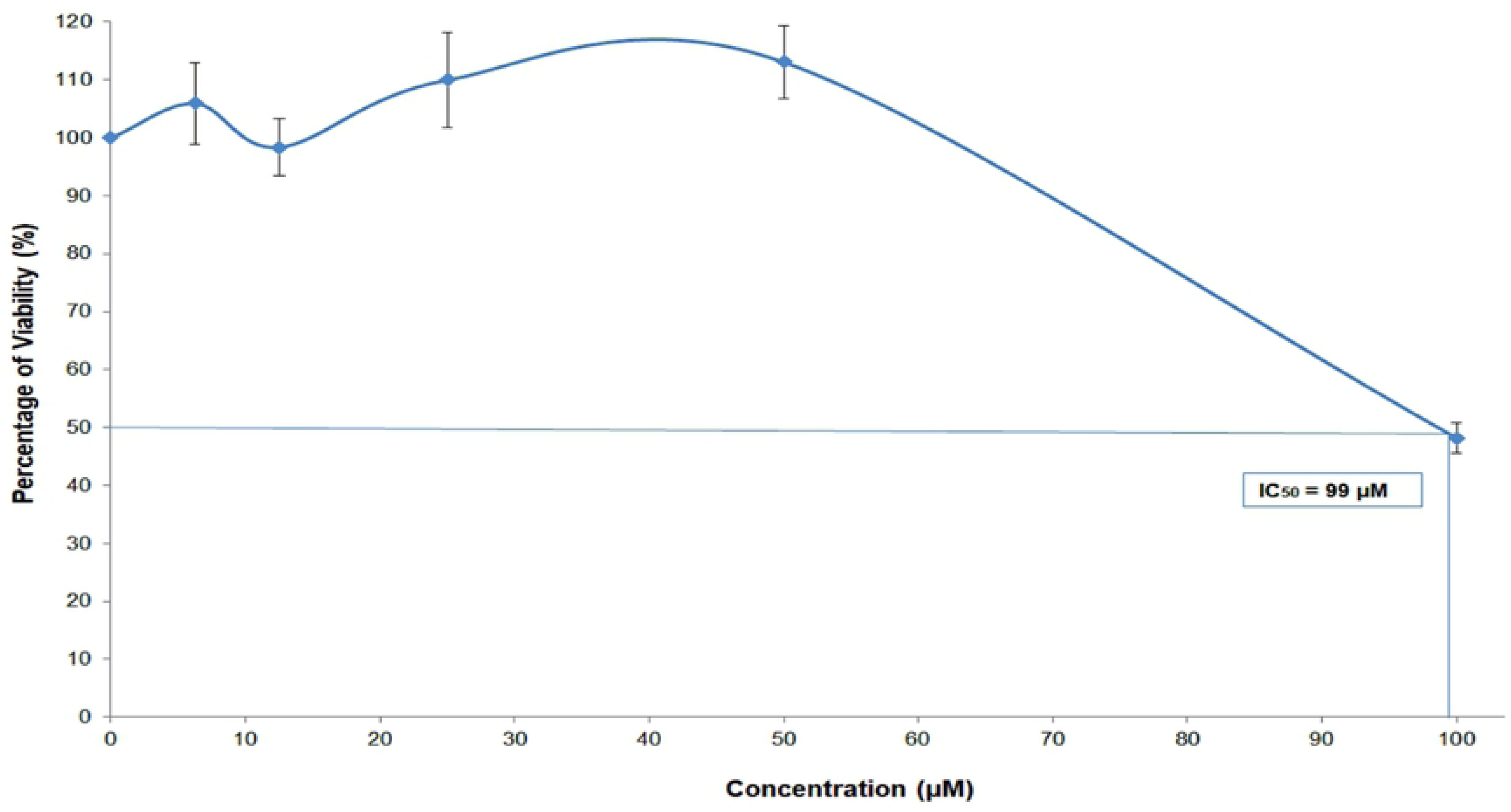
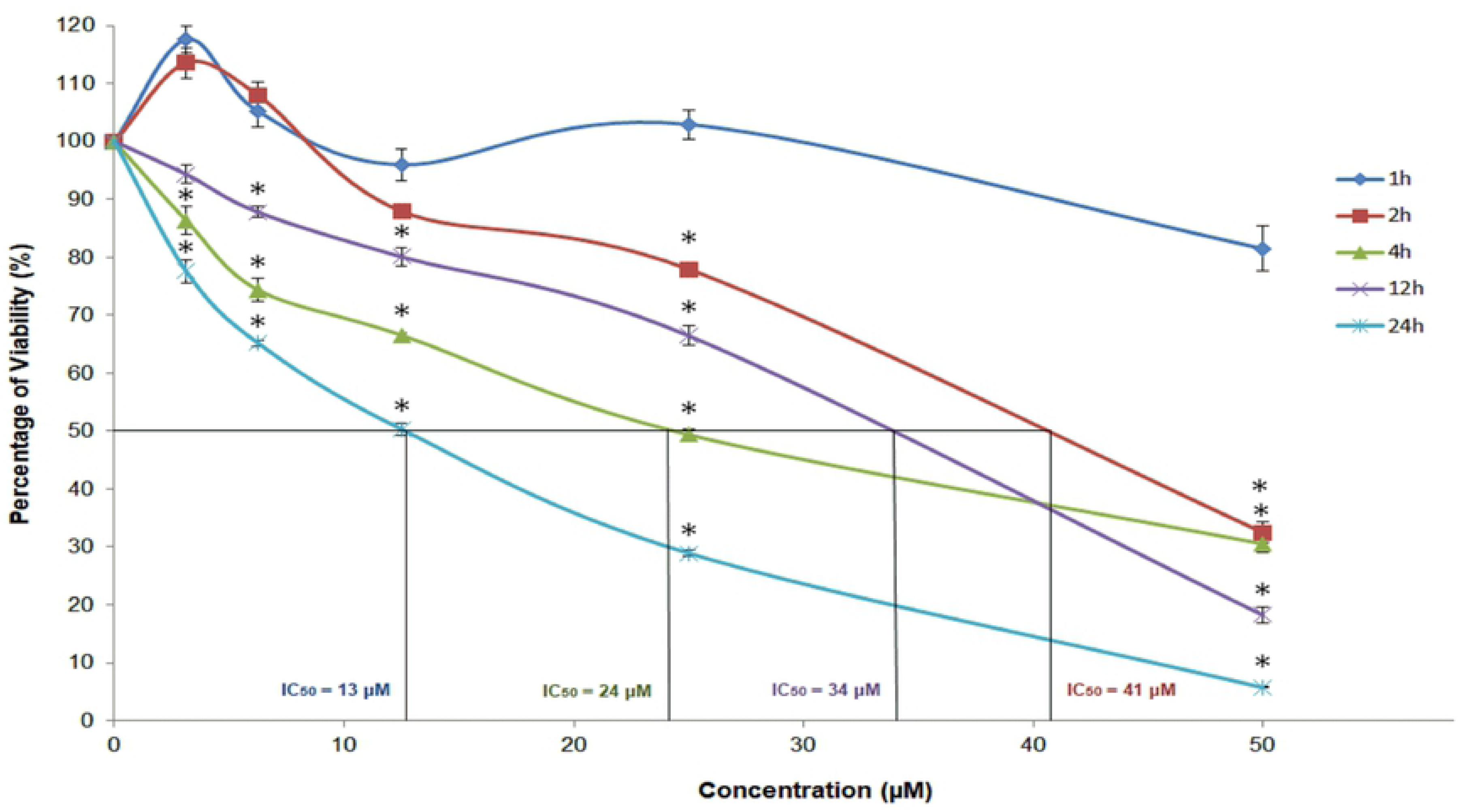
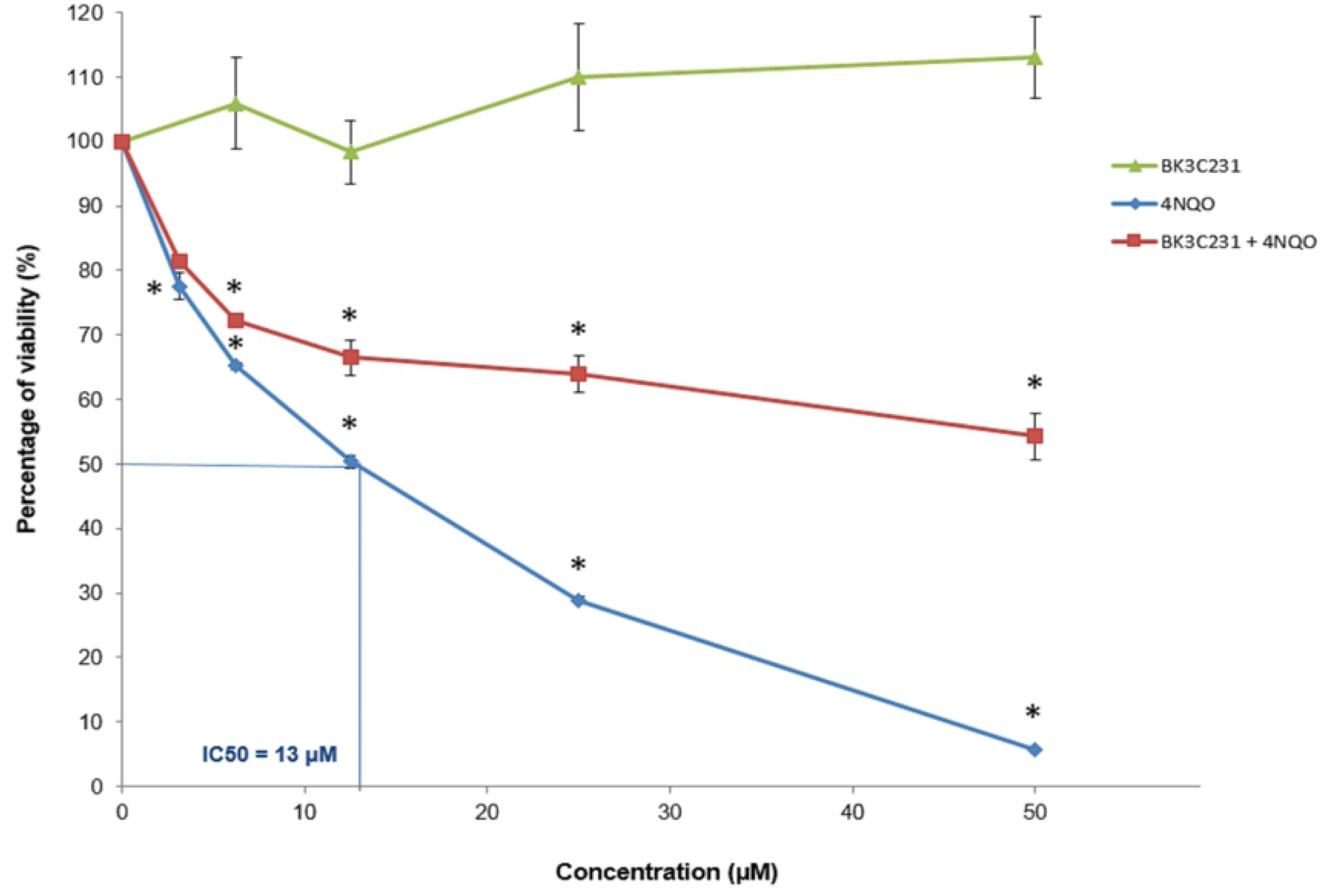
Effect of BK3C231 and 4NQO on the viability of CCD-18Co cells as assessed by MTT assay. **(A)** Cells were treated with BK3C231 from 6.25 µM till 100 µM for 24h. An IC_50_ value of 99 µM was observed. **(B)** Cells were treated with 4NQO from 3.125 µM till 50 µM for 1h (no IC_50_ value), 2h (IC_50_ value was 41 μM), 4h (IC_50_ value was 24 μM), 12h (IC_50_ value was 34 μM) and 24h (IC_50_ value was 13 μM). **(C)** Cells were pretreated with BK3C231 at 50 μM for 2h prior to 4NQO treatment from 3.125 μM till 50 μM for subsequent 22h (no IC_50_ value) in comparison to BK3C231-treated (no IC_50_ value) and 4NQO-treated cells (IC_50_ value was 13 μM). Each data point was obtained from three independent experimental replicates and expressed as mean ± SEM of percentage of cell viability. * p<0.05 against negative control.

### 3.2 BK3C231 protection against 4NQO-induced DNA microlesions

Significant DNA damage as indicated by the comet tail which represents DNA strand breaks can be observed in cells treated only with 4NQO. Untreated control cells and BK3C231-treated cells showed intact round nuclear DNA and no DNA strand break was observed at all treated concentrations (Fig 3A). There was also a decrease in comet tail in cells pretreated with BK3C231 when compared with that of cells treated only with 4NQO (Fig 3B). This was further confirmed by quantification of tail moments obtained from comet scoring. Tail moment increased significantly up to 48-fold in 4NQO-treated cells at 28.79 ± 1.02 (p<0.05) over control and BK3C231-treated cells ranging from 0.59 ± 0.11 to 0.68 ± 0.06 (Fig 4A). On the other hand, BK3C231 pretreatment showed a 0.8-fold decrease of 4NQO-induced DNA strand breaks in a concentration-dependent manner, significantly at 50 µM with a tail moment value of 7.21 ± 0.34 (p<0.05) (Fig 4B).

**Fig 3.**
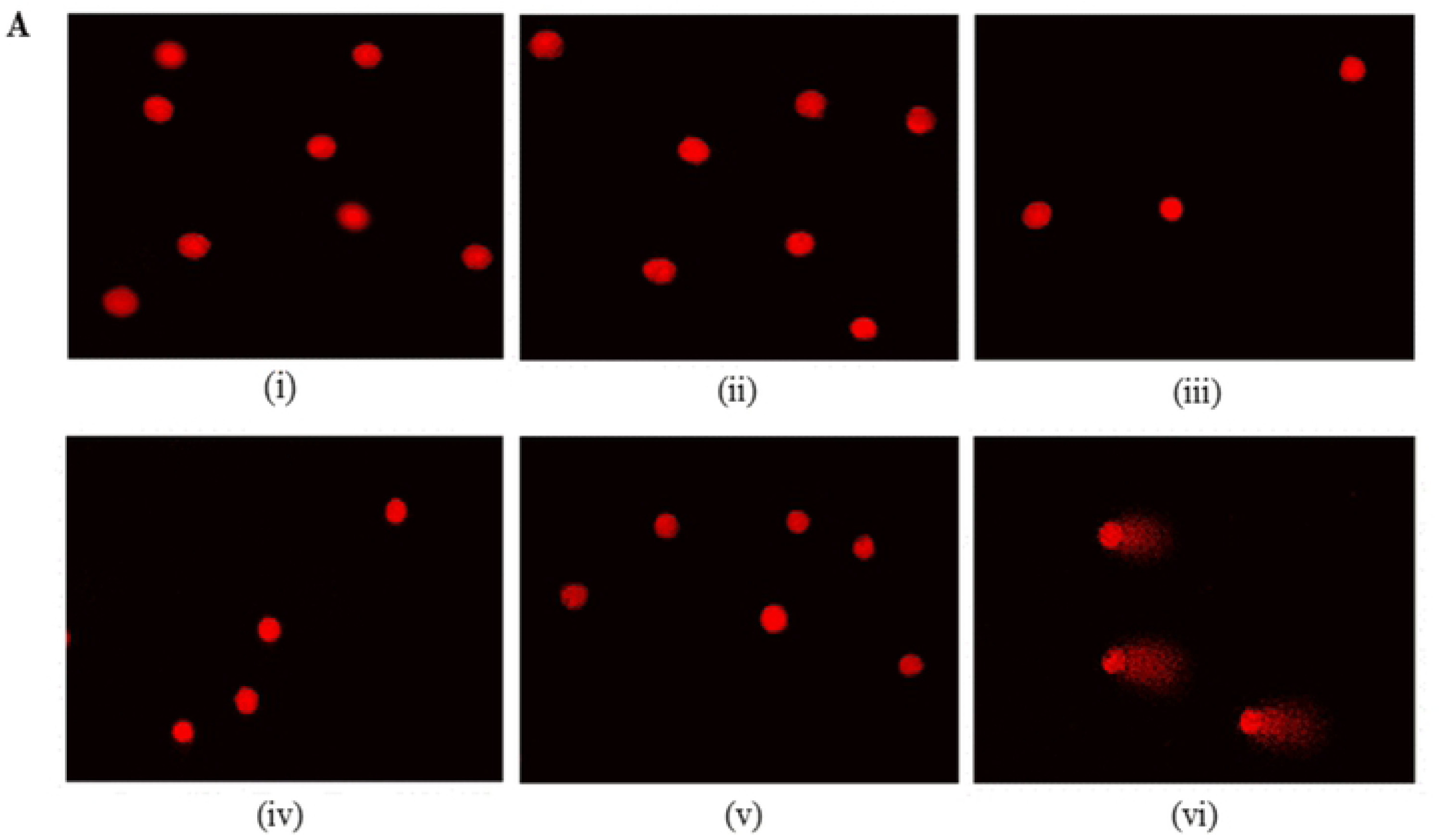
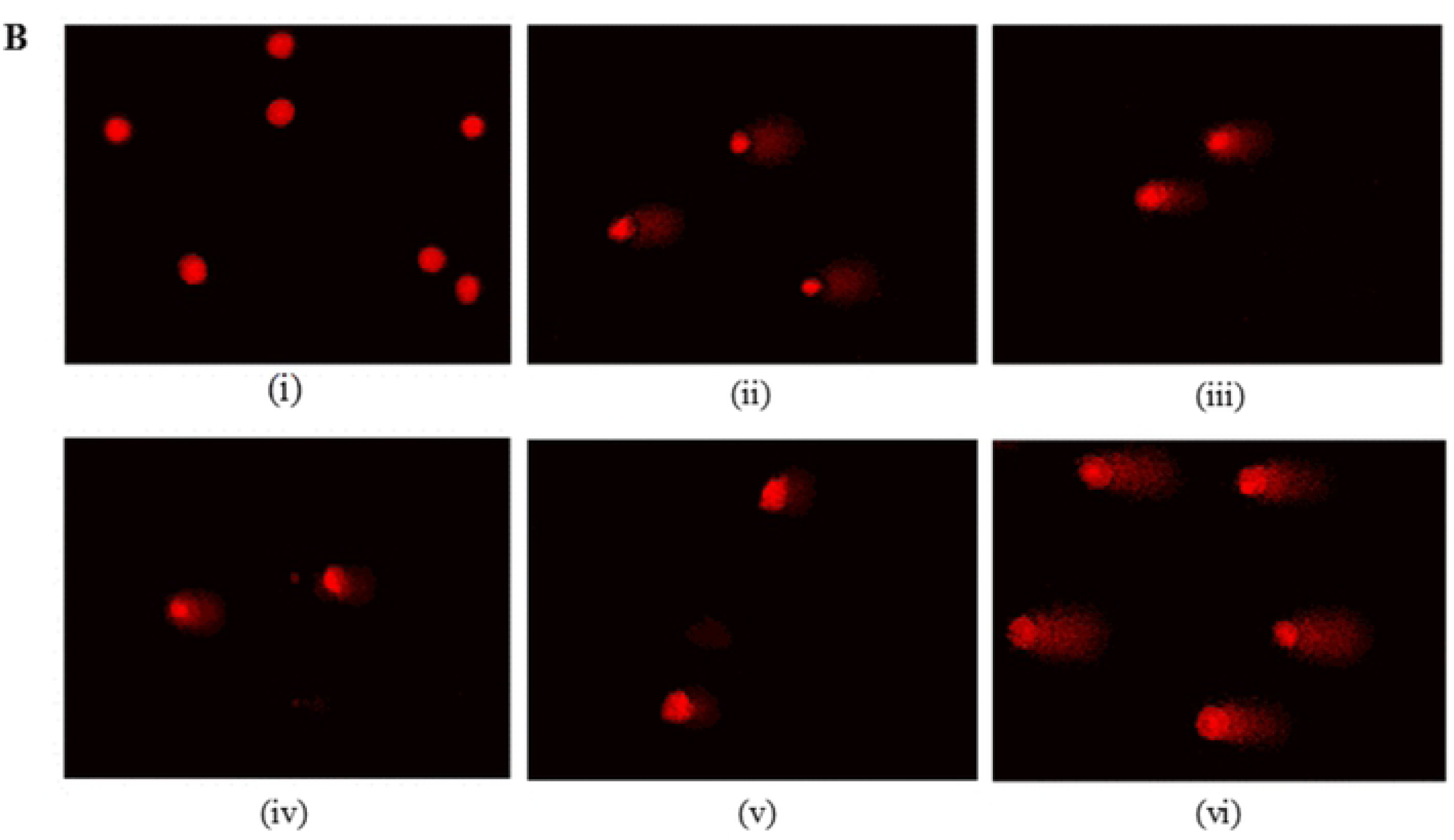
DNA microlesion assessment in CCD-18Co cells using Alkaline Comet assay. (A) Fluorescence microscopic images with EtBr staining of untreated cells (i), cells treated with BK3C231 at 6.25 µM (ii), 12.5 µM (iii), 25 µM (iv) and 50 µM (v) for 24h and cells treated with 4NQO at 1 µM for 1h (vi). **(B)** Fluorescence microscopic images of untreated cells (i), cells treated with BK3C231 at 6.25 µM (ii), 12.5 µM (iii), 25 µM (iv) and 50 µM (v) for 2h prior to 4NQO induction at 1 µM for 1h and cells treated with 4NQO at 1 µM for 1h (vi). Each data represents at least three independent experimental replicates.

**Fig 4.**
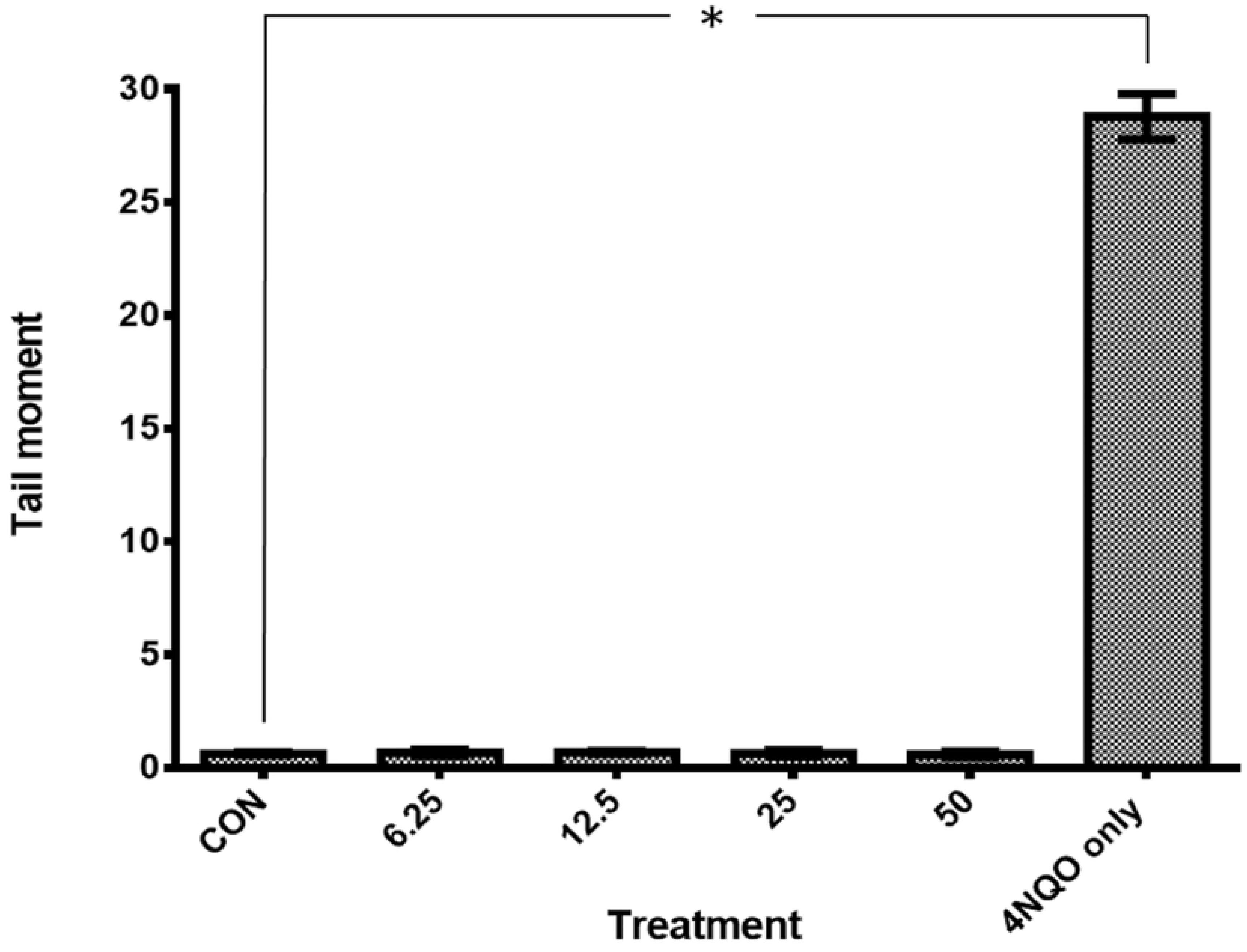
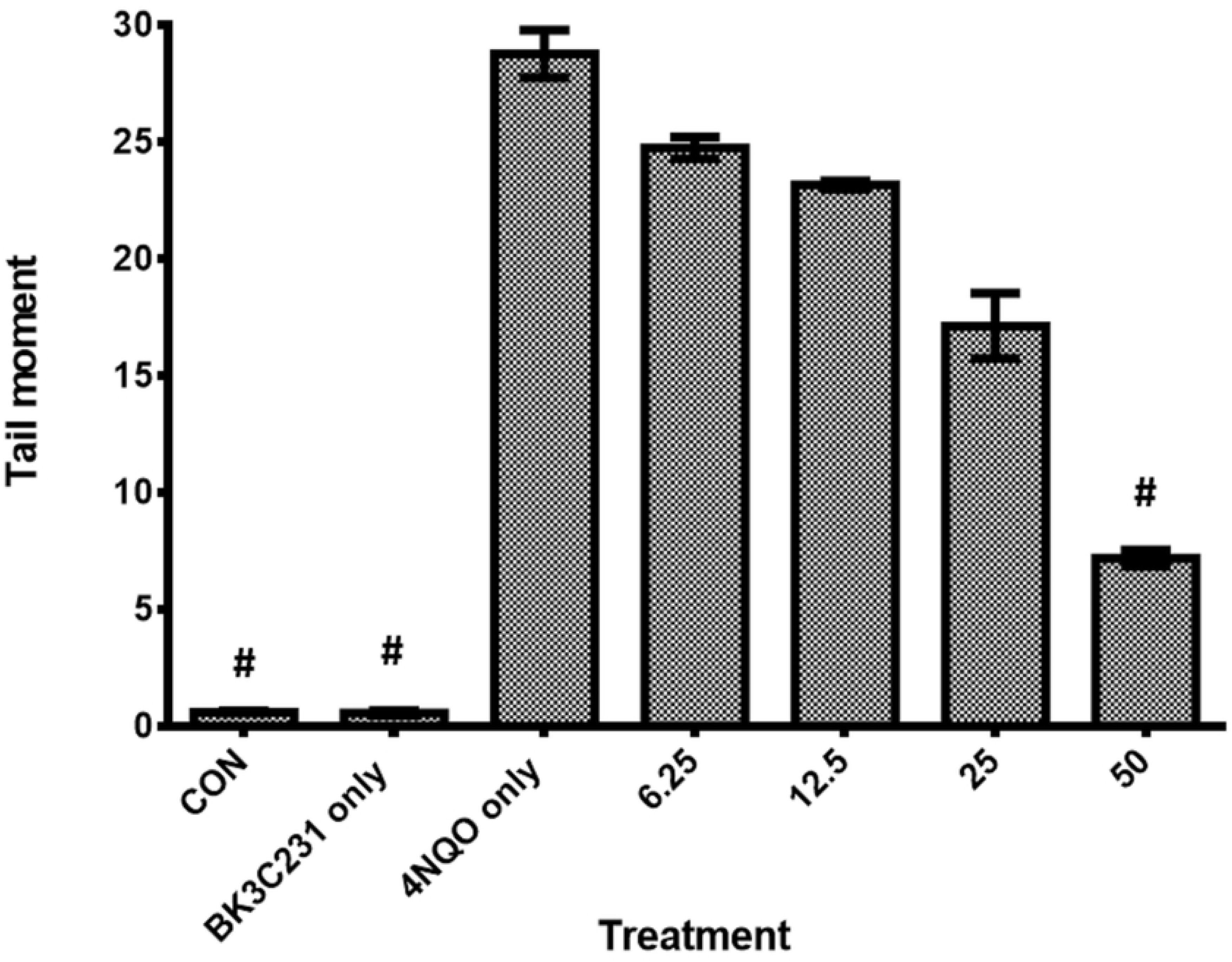
Tail moments obtained from comet scoring in CCD-18Co cells. (A) Screening for DNA damage expressed as tail moment in cells treated respectively with BK3C231 from 6.25 µM till 50 µM for 24h and 4NQO at 1 µM for 1h. **(B)** Cells were pretreated with BK3C231 from 6.25 µM till 50 µM for 2h prior to 4NQO induction at 1 µM for 1h. Each data point was obtained from three independent experimental replicates and expressed as mean ± SEM of tail moment. * p<0.05 against negative control, CON (A) and # p<0.05 against positive control, 4NQO only (B).

### 3.3 Inhibition of 4NQO-induced DNA macrolesions by BK3C231

The protective role of BK3C231 against 4NQO-induced micronucleus formation was assessed using CBMN assay. In untreated control cells, a micronucleus frequency level as low as 0.23 ± 0.03 was observed. Cells treated with 4NQO significantly demonstrated up to 93-fold increase in frequency of micronucleus in binucleated cells at 21.56 ± 1.36 (p<0.05). However, pretreatment of cells with BK3C231 was shown to cause a maximum of 0.8 fold decrease of 4NQO-induced micronucleus formation in a concentration-dependent manner, significantly at 25 µM with a frequency level of 6.58 ± 0.52 and 50 µM with a frequency level of 3.80 ± 0.47 (p<0.05) (Fig 5B). In addition, the NDI values measured in control, 4NQO-treated cells and BK3C231-treated cells were 1.78 ± 0.01, 1.68 ± 0.03 and 1.79 ± 0.01 respectively. As for cells pretreated with BK3C231 prior to 4NQO induction, the average NDI value measured was 1.72 ± 0.01 (data not shown). All NDI values obtained in this assay indicated normal cell proliferation [39].

**Fig 5.**
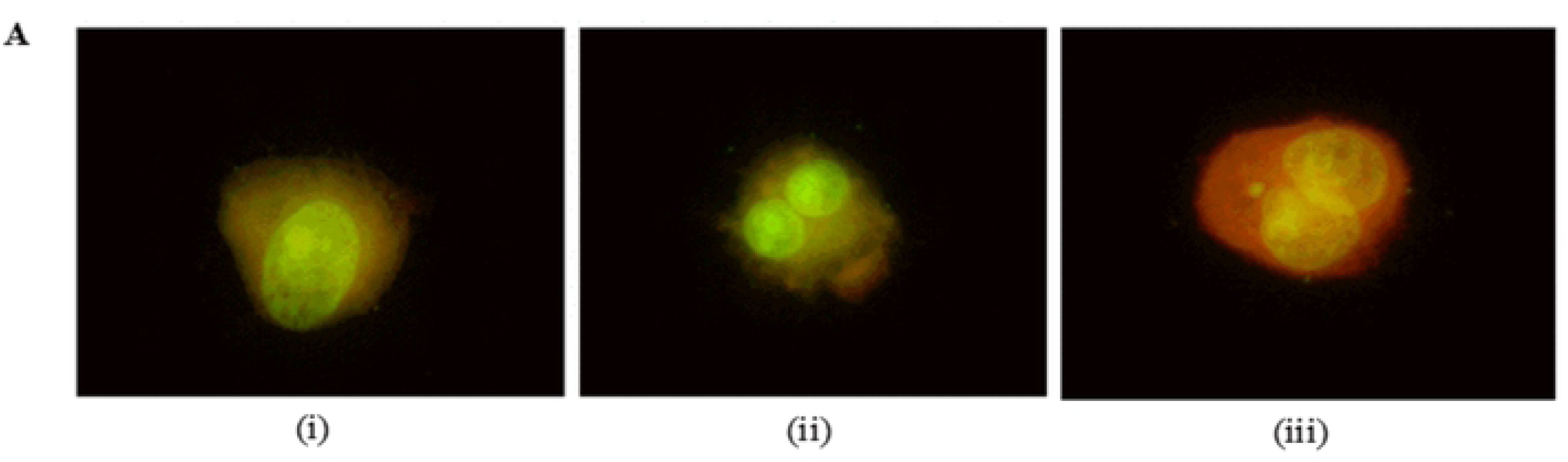
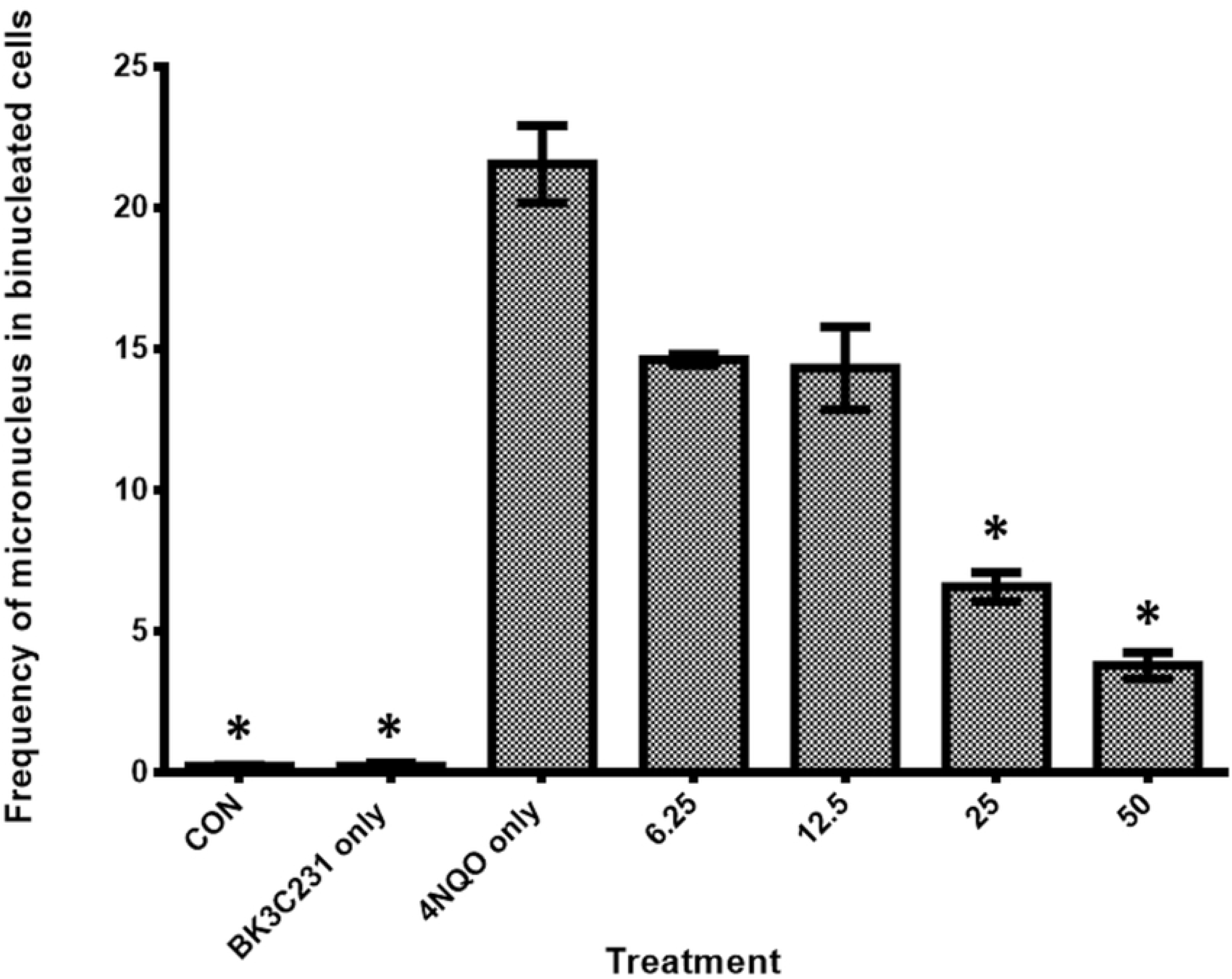
DNA macrolesion assessment in CCD-18Co cells using CBMN assay. (A) Fluorescence microscopic images with acridine orange staining of mononucleated cell (i), binucleated cell (ii) and binucleated cell with micronucleus (iii). Cellular nucleus was stained green while cytoplasm was stained orange in this assay. **(B)** Cells were pretreated with BK3C231 from 6.25 µM till 50 µM for 2h prior to 4NQO induction at 1 µM for 2h. Each data point was obtained from three independent experimental replicates and expressed as mean ± SEM of frequency of micronucleus in binucleated cells. * p<0.05 against positive control, 4NQO only.

### 3.4 Cytoprotective role of BK3C231 in 4NQO-induced loss of mitochondrial membrane potential (ΔΨm)

The cytoprotective role of BK3C231 was further investigated at the mitochondrial level through flow cytometric assessment of ΔΨm loss using TMRE staining. Significant loss of ΔΨm (p<0.05) as indicated by a 1.2-fold increase of TMRE-negative cells from 15.63 ± 1.09 % in control cells to 34.77 ± 1.29 % in 4NQO-treated cells was observed. However, BK3C231 pretreatment was shown to reduce the amount of TMRE-negative cells significantly to 22.13 ± 2.51 % (p<0.05) at 50 µM, thereby protecting the cells against 4NQO-induced ΔΨm loss (Fig 6).

**Fig 6.**
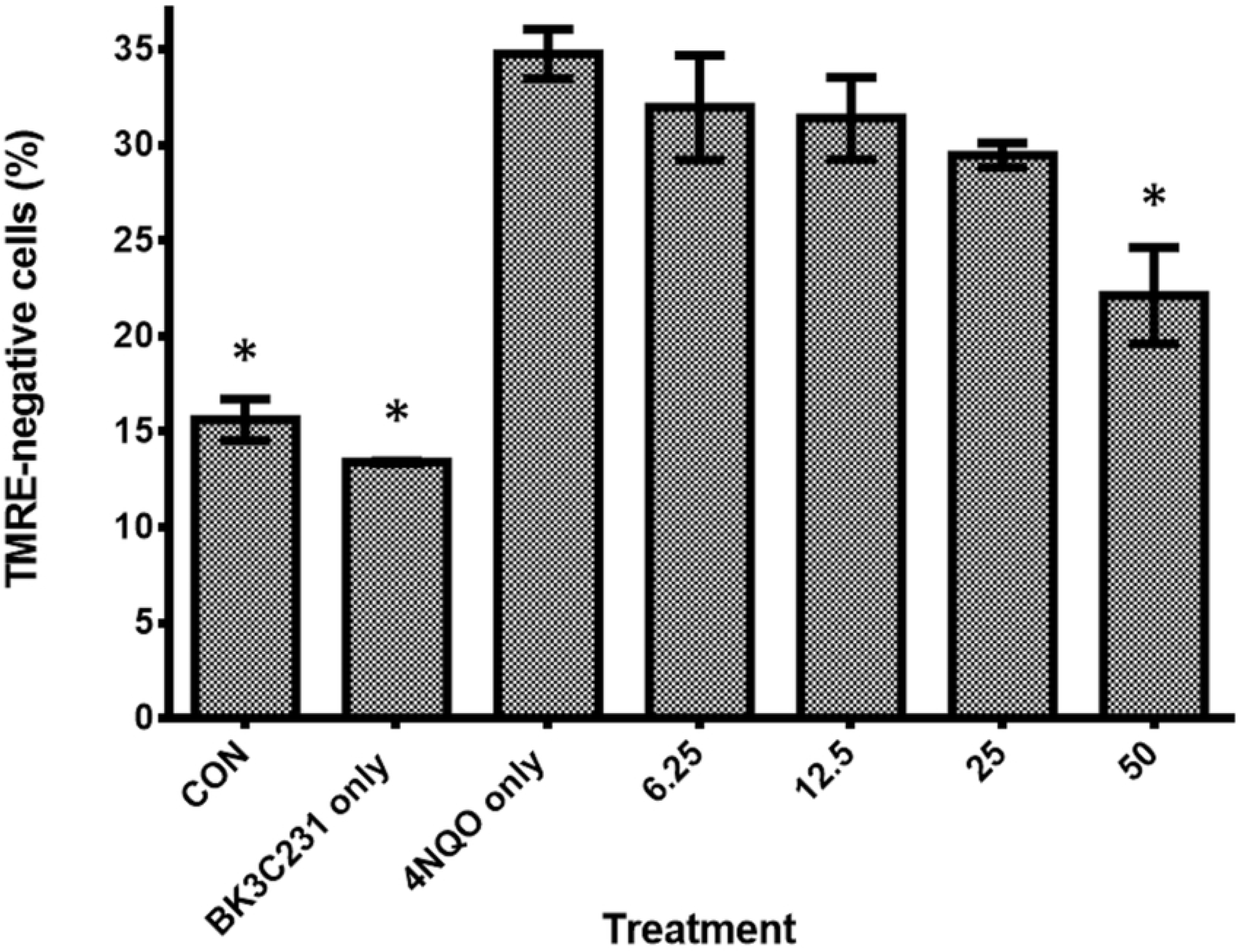
Flow cytometric assessment of ΔΨm level using TMRE staining. Cells were pretreated with BK3C231 from 6.25 µM till 50 µM for 2h prior to 4NQO induction at 1 µM for 2h. Each data point was obtained from three independent experimental replicates and expressed as mean ± SEM of TMRE-negative cells (%). * p<0.05 against positive control, 4NQO only.

### 3.5 Suppression of 4NQO-induced cardiolipin loss by BK3C231

In a bid to further establish the protective role of BK3C231 in mitochondria, cardiolipin level was assessed through flow cytometric analysis using NAO staining. Our study demonstrated significant cardiolipin loss (p<0.05) as indicated by a 2.8-fold increase of NAO-negative cells from 5.07 ± 0.52 % in control cells to 19.33 ± 0.94 % in 4NQO-treated cells. However, BK3C231 pretreatment was shown to induce up to 0.4-fold decrease of 4NQO-induced cardiolipin loss in a concentration-dependent manner (Fig 7).

**Fig 7.**
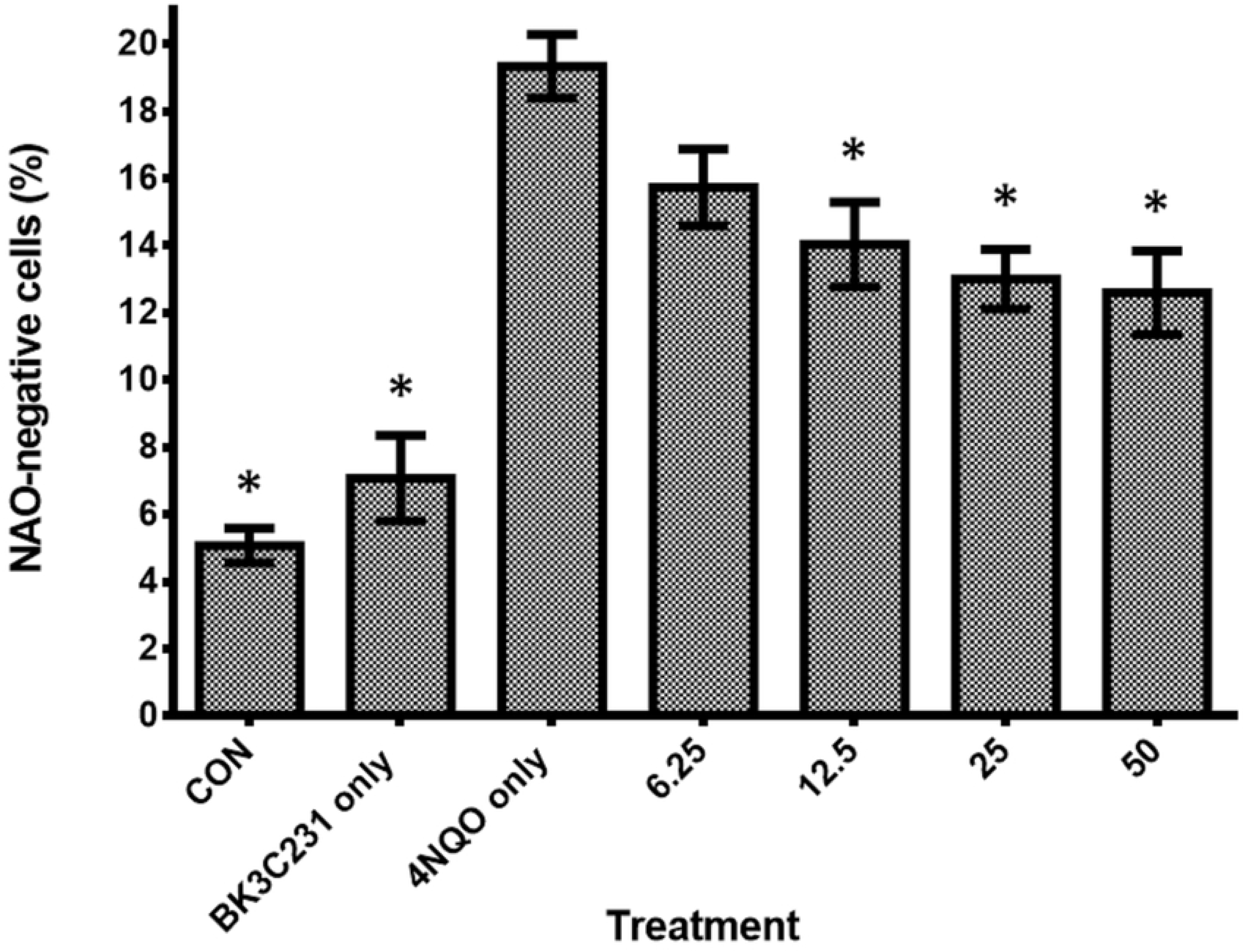
Flow cytometric assessment of cardiolipin level using NAO staining. Cells were pretreated with BK3C231 from 6.25 µM till 50 µM for 2h prior to 4NQO induction at 1 µM for 2h. Each data point was obtained from three independent experimental replicates and expressed as mean ± SEM of NAO-negative cells (%).* p<0.05 against positive control, 4NQO only.

### 3.6 4NQO-induced DNA and mitochondrial damages independent of ROS production

Flow cytrometric assessment of ROS namely superoxide and hydrogen peroxide levels using HE and DCFH-DA staining was performed to determine the role of ROS in 4NQO-induced DNA and mitochondrial damages. Interestingly, as shown in Fig 8A-B, there were no inductions of superoxide and hydrogen peroxide levels in 4NQO-treated cells as compared to control cells. Hydroquinone (HQ), which was used as positive control, had significantly increased ROS level in CCD-18Co cells (p<0.05). This suggested that ROS was not involved in DNA and mitochondrial damages caused by 4NQO.

**Fig 8.**
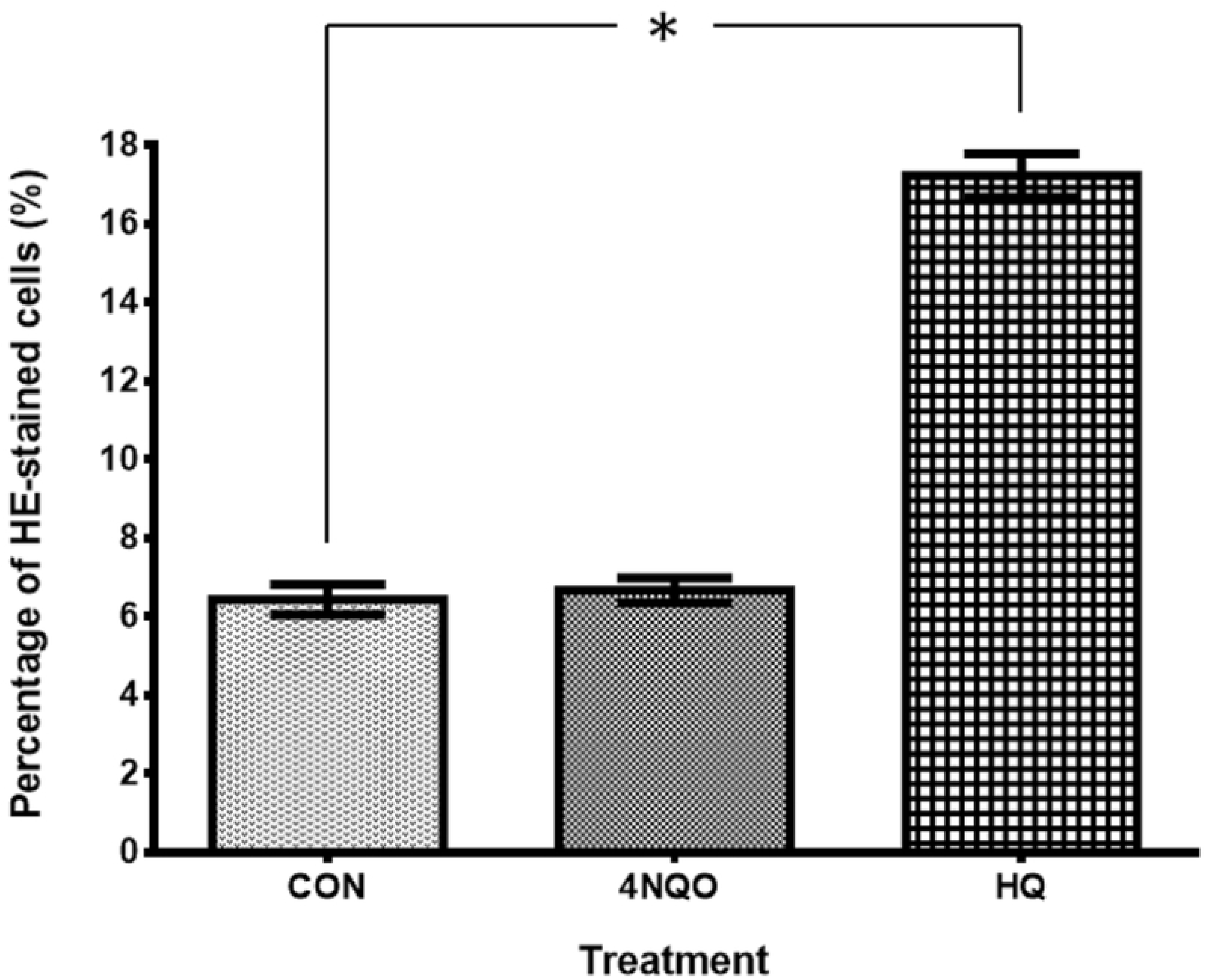
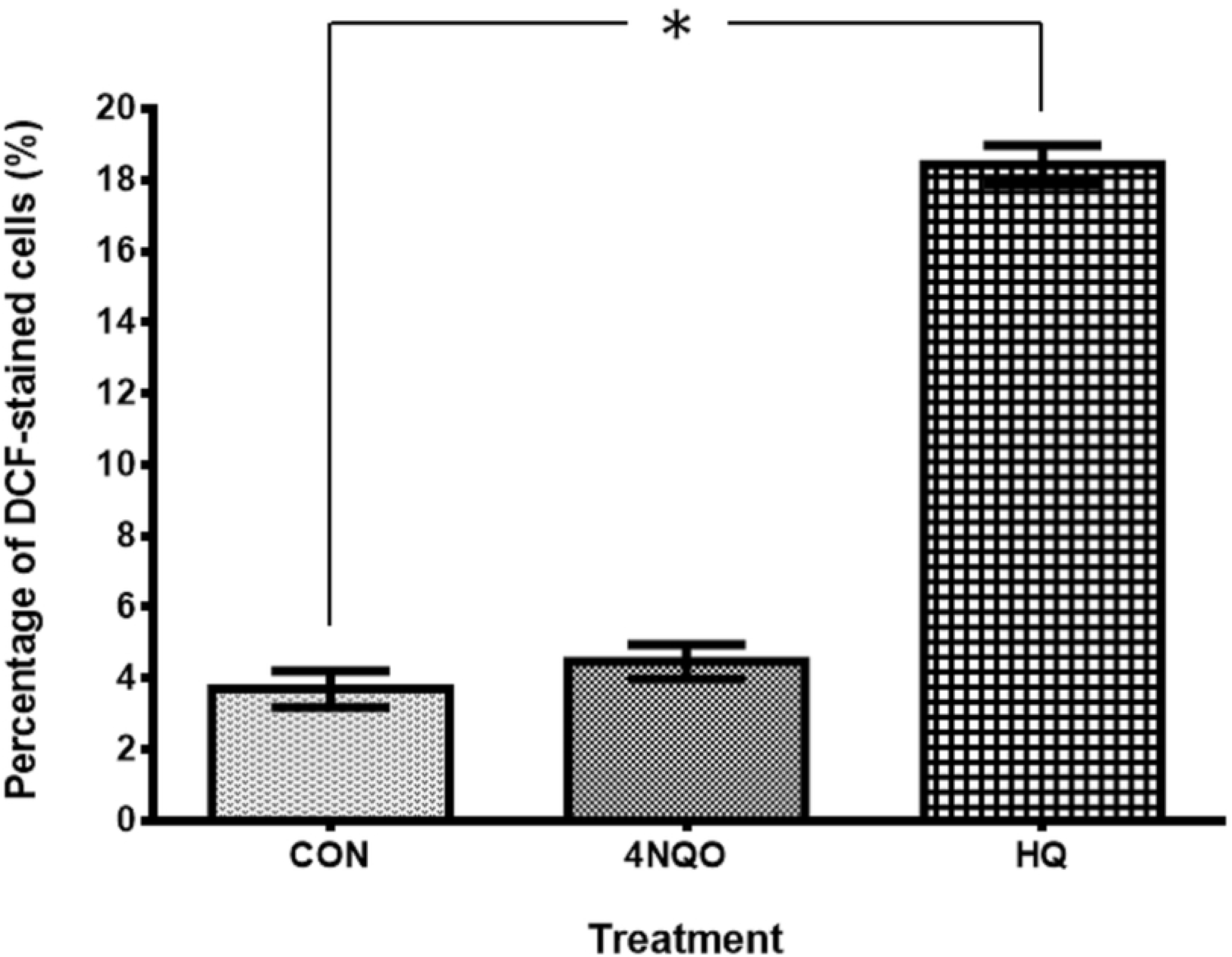
Flow cytometric assessment of superoxide level using HE staining (A) and hydrogen peroxide level using DCFH-DA staining (B). Cells were treated with 4NQO at 1 µM for 1h. HQ treatment at 50 µM for 2h was used as positive control in this assay. Each data point was obtained from three independent experimental replicates and expressed as mean ± SEM of percentage of HE- or DCF-stained cells. * p<0.05 against negative control, CON.

### 3.7 Inhibition of 4NQO-induced nitrosative stress by BK3C231

Intracellular nitric oxide (NO) level was assessed using BD Pharmingen™ Orange NO Probe staining whereas extracellular NO level was assessed using Griess assay to determine the involvement of RNS in 4NQO-induced DNA and mitochondrial damages. Our study demonstrated a significant 0.98-fold increase of intracellular NO level, 15.7 ± 0.19 % and 2.4-fold increase of extracellular NO level, 5.15 ± 0.17 µM (p<0.05) in 4NQO-treated cells over control cells, at 7.63 ± 0.19 % and 1.51 ± 0.26 µM respectively, thereby demonstrating the involvement of NO in 4NQO-induced DNA and mitochondrial damages. Moreover, BK3C231 significantly inhibited 4NQO-induced NO production from as early as 2 hours up till 24 hours of pretreatment (p<0.05) (Fig 9A,B). In addition to that, antioxidant GSH level was assessed using Ellman’s reagent. 4NQO-treated cells showed a reduced GSH level at 194.70 ± 23.83 nmol/mg as compared to untreated control cells at 245.96 ± 12.44 nmol/mg (Fig 9C). Overall, the simultaneous increase in NO level and decrease in GSH level by 4NQO further confirmed the involvement of nitrosative stress in 4NQO-induced DNA and mitochondrial damages. However, no induction of GSH level was observed in cells pretreated with BK3C231 for 2 hours, 4 hours, 6 hours and 12 hours. BK3C231 was only able to significantly increase GSH level, 313.97 ± 27.83 nmol/mg (p<0.05) at 24 hours of pretreatment as compared to that of 4NQO-treated cells (Fig 9C). This suggested that BK3C231 inhibited 4NQO-induced nitrosative stress through early reduction of NO production and late induction of GSH level in CCD-18Co cells.

**Fig 9.**
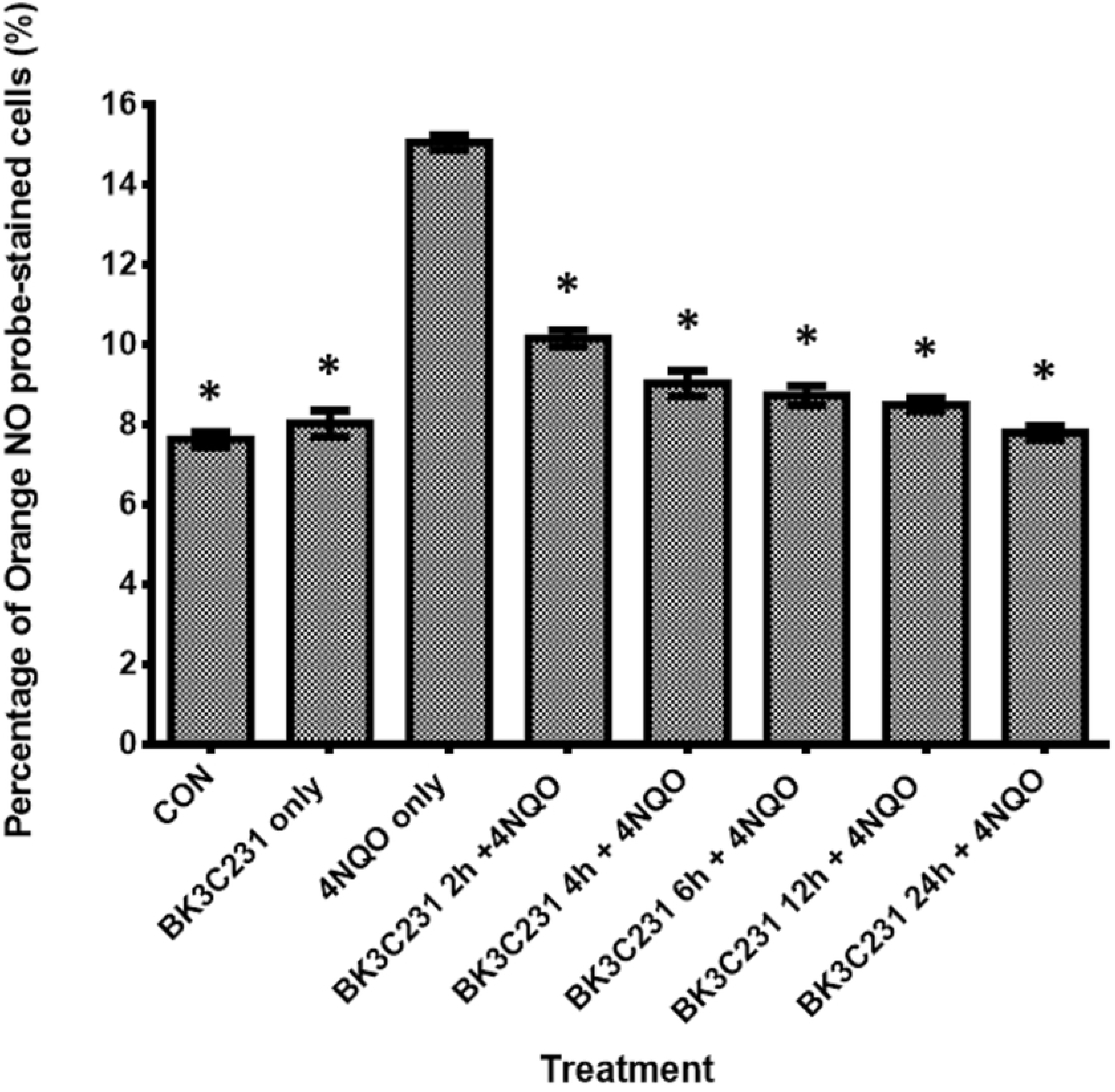
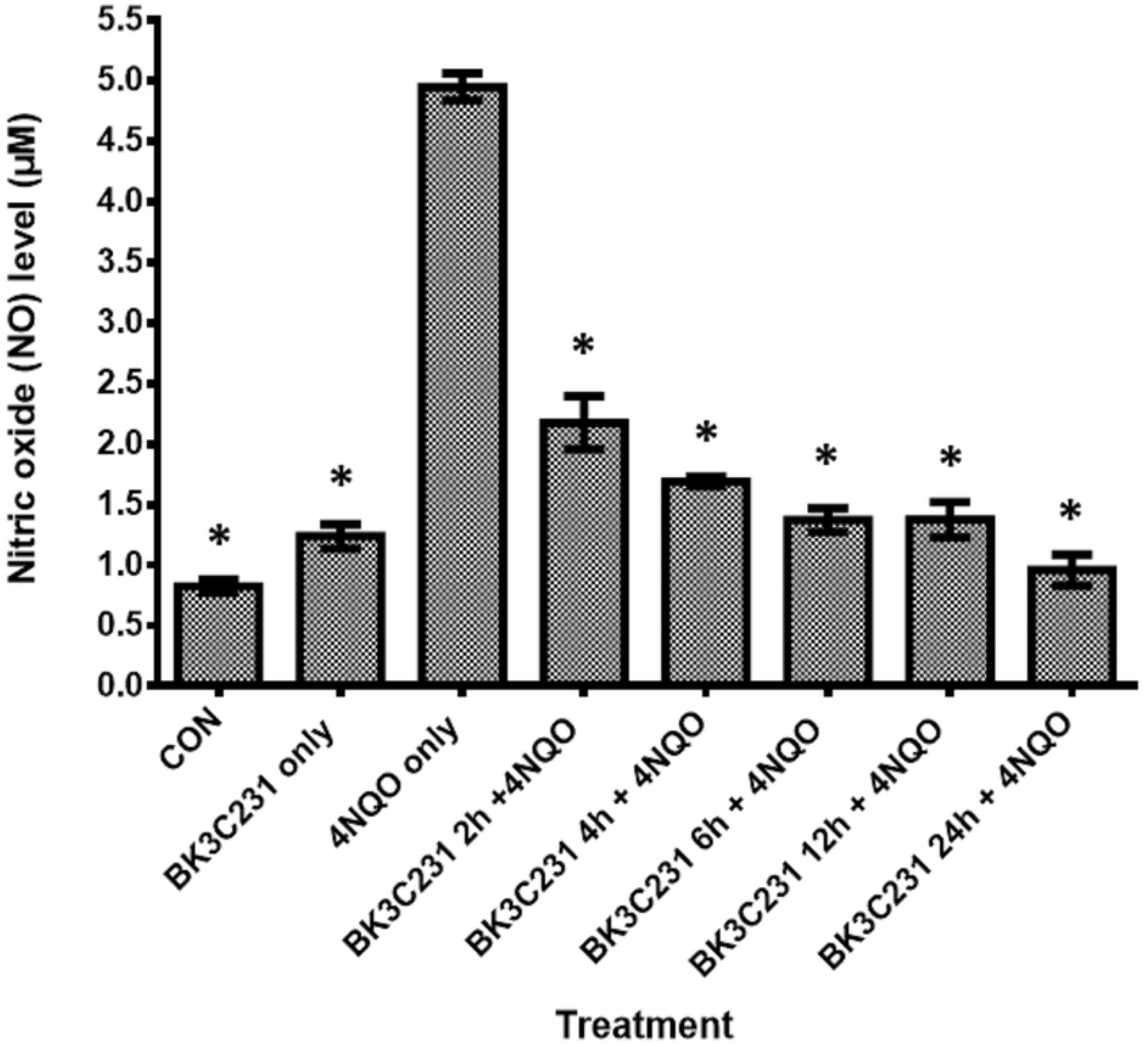
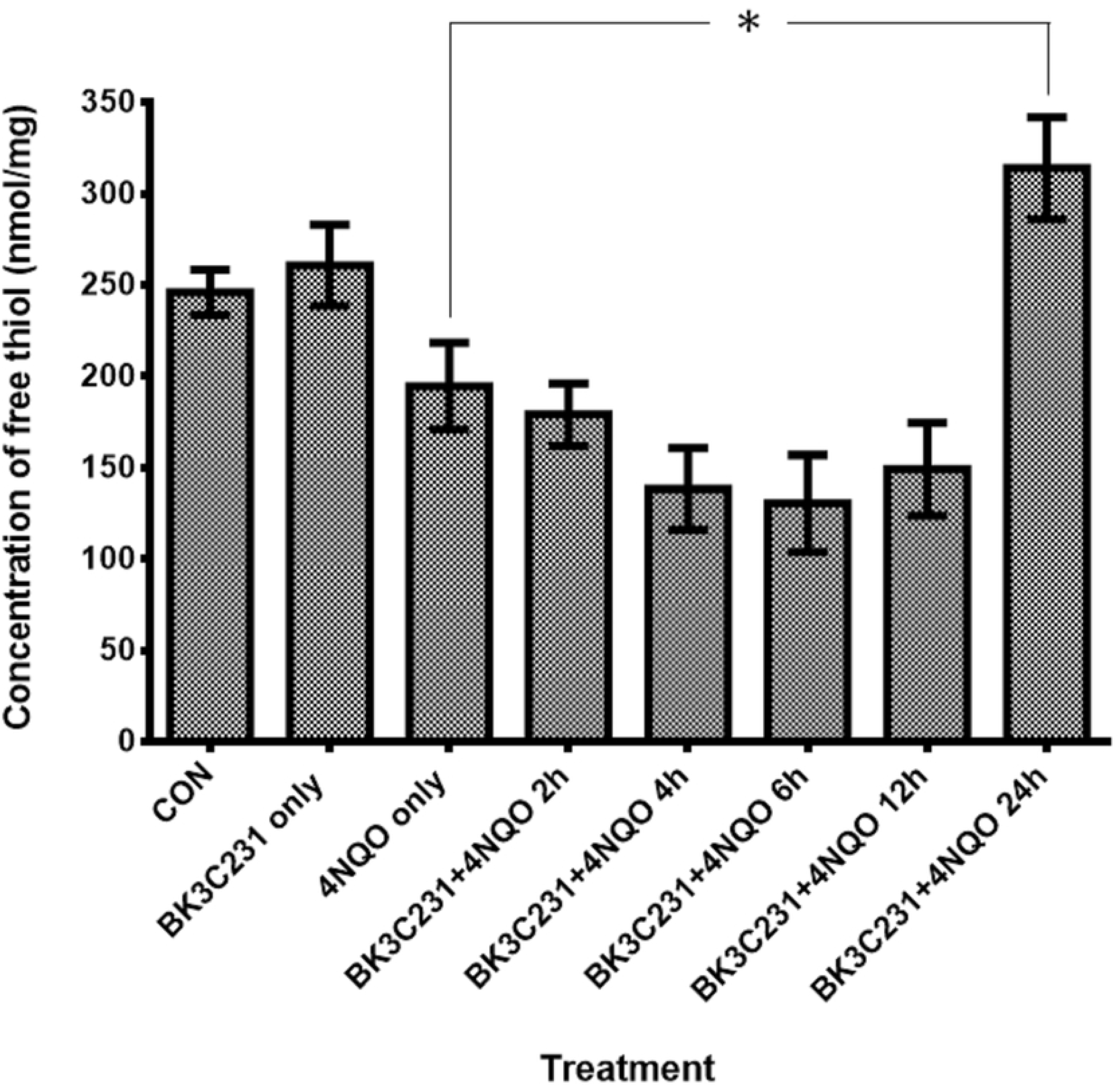
Nitrosative stress assessment through determination of intracellular NO level using Orange NO probe staining (A), extracellular NO level using Griess assay (B) and GSH level using Ellman’s reagent (C) in CCD-18Co cells. Cells were pretreated with BK3C231 at 50 µM for 2h, 4h, 6h, 12h and 24h prior to 4NQO induction at 1 µM for 1h. Each data point was obtained from three independent experimental replicates and expressed as mean ± SEM of NO level and concentration of free thiol. * p<0.05 against positive control, 4NQO only.

## 4. Discussion

Epidemiological studies have shown that consumption of fruits particularly rich in stilbenes led to a reduced risk of colorectal cancer, which is one of the most commonly diagnosed cancer worldwide [40, 41]. Furthermore, cytoprotection of DNA and mitochondrial function limits the occurrence of cancer. Since DNA is the repository of hereditary material and genetic information in every living cell, the maintenance of its stability is pivotal as unrepaired DNA damages caused by diverse assaults from the environment, nutrition and natural cellular processes lead to cancer [42, 43]. As for mitochondria, impairments and alterations of mitochondrial structure and functions, including morphology and redox potential, are associated with cancer transformation and have been frequently reported in human cancers [44–46]. In agreement with this, our study showed that BK3C231 was able to inhibit 4NQO-induced cytotoxicity as well as protect against DNA damage and mitochondrial dysfunction in the normal human colon fibroblast CCD-18Co cell line.

Firstly, we sought to understand the carcinogenic actions of 4NQO. Studies have demonstrated that 4NQO elicited carcinogenicity through its proximate carcinogenic metabolite namely 4-hydroxyaminoquinoline 1-oxide (4HAQO) produced by the enzymatic four-electron reduction of 4NQO’s nitro group [47, 48]. Being a potent chemical carcinogen and as a UV-mimetic agent, 4NQO is often used as positive control in various genototoxicity studies due to its well characterized metabolic processes [49]. Though a study by Brüsehafer et al. [50] reported that 4NQO predominantly induces mutagenicity more than clastogenicity and that the latter depends on cell types, our study has proved that 4NQO significantly caused DNA damage via DNA strand breaks and chromosomal damage via micronucleus formation. Also, our study was in agreement with previous studies which demonstrated that 4NQO caused damage to mitochondrial membrane as characterized by loss of mitrochondrial membrane potential (ΔΨm) and cardiolipin [51].

As 4HAQO’s carcinogenic effect is mainly based on DNA adduct formation, our study investigated 4NQO’s other carcinogenic mechanism of action through generation of ROS and RNS and its involvement in the cytoprotective role of BK3C231 [52–54]. Interestingly, our study which revealed no superoxide and hydrogen peroxide production by 4NQO at 1 µM for 1 hour in CCD-18Co cells contradicts the study by Arima et al. [37] which reported ROS formation in human primary skin fibroblast by 4NQO using the same treatment concentration and timepoint. The discrepancy is likely due to the difference in the origin of fibroblast used. Hence, our study is the first to elucidate such findings on 4NQO mechanism which has never been shown in other studies thus far.

In addition, our study demonstrated an increased NO level and a depleted GSH level by 4NQO. This is possibly due to formation of 4NQO-GSH conjugates leading to generation of nitrite, a stable end product of NO, which inactivated γ-glutamylcysteine synthase and therefore suppressed intracellular synthesis of GSH [37,54–56]. Our data was also in agreement with previous studies that NO could be the main culprit in 4NQO-induced DNA and mitochondrial damages in CCD-18Co cells as NO has been demonstrated to induce genotoxicity and damage to mitochondria via multiple mechanisms directly or indirectly [57, 58]. NO also plays an important role in tumour biology and overproduction of NO can promote tumour growth [59]. Moreover, the concurrent increase in NO level and decrease in GSH level postulates the occurrence of nitrosative stress which may contribute to 4NQO-induced DNA and mitochondrial damages [60, 61].

More importantly, BK3C231 was shown in our study to protect against 4NQO-induced DNA and mitochondrial damages by decreasing DNA strand breaks and micronucleus formation as well as reducing loss of mitochondrial membrane potential (ΔΨm) and cardiolipin. Our study further revealed that BK3C231 exerted these cytoprotective effects in CCD-18Co cells by suppressing 4NQO-induced nitrosative stress through reduction in NO level and late upregulation of GSH level. The role of stilbene derivatives as potential antioxidants has been a conventional fact proven by many studies such as resveratrol, a well-known stilbenoid, attenuated nitrosative stress in small intestine of rats [62]. Piceatannol and isorhapontigenin, which are natural occurring stilbenes, have also been demonstrated to scavenge NO and nitrogen dioxide (NO_2_) radicals as well as increasing GSH/GSSG ratio [63, 64].

Kee et al. [36] has reported chemopreventive activity of BK3C231 involving upregulation of detoxifying enzyme NQO1 due to the presence of methoxy group and furan carboxamide. Therefore, it is possible that the depletion of NO level is mediated directly by BK3C231 most likely due to the presence of methoxy group that enables BK3C231 to act as free radical scavenger which donates electron to scavenge NO. Besides that, the upstream Keap1-Nrf2 signaling pathway, which is a major regulator of phase II detoxification and cytoprotective genes, is postulated to be involved through upregulation of detoxifying enzymes which may lead to NO suppression. Stilbene derivatives particularly resveratrol have been playing substantial role in activation of Nrf2-related gene transcription which induces expression of cytoprotective enzymes such as NQO1, glutathione S-transferase (GST), glutamate-cysteine ligase catalytic subunit (GCLC) and heme oxygenase-1 (HO-1) thus leading to protection against cancer [65]. Therefore, our study warrants further investigation on the role of BK3C231 in the Keap1-Nrf2 pathway.

## 5. Conclusion

In conclusion, this study has provided a better insight into 4NQO-induced carcinogenicity in CCD-18Co cells. Our findings also served as a stepping stone for further elucidation of BK3C231 chemopreventive potential against both genetic and epigenetic bases of cancer development. Through these findings, we aim to design BK3C231 as an ideal chemopreventive agent in hopes of reducing the gap between understanding molecular mechanism occurring in cancer carcinogenesis and instigating successful adoption of chemoprevention.

**Fig 10.**
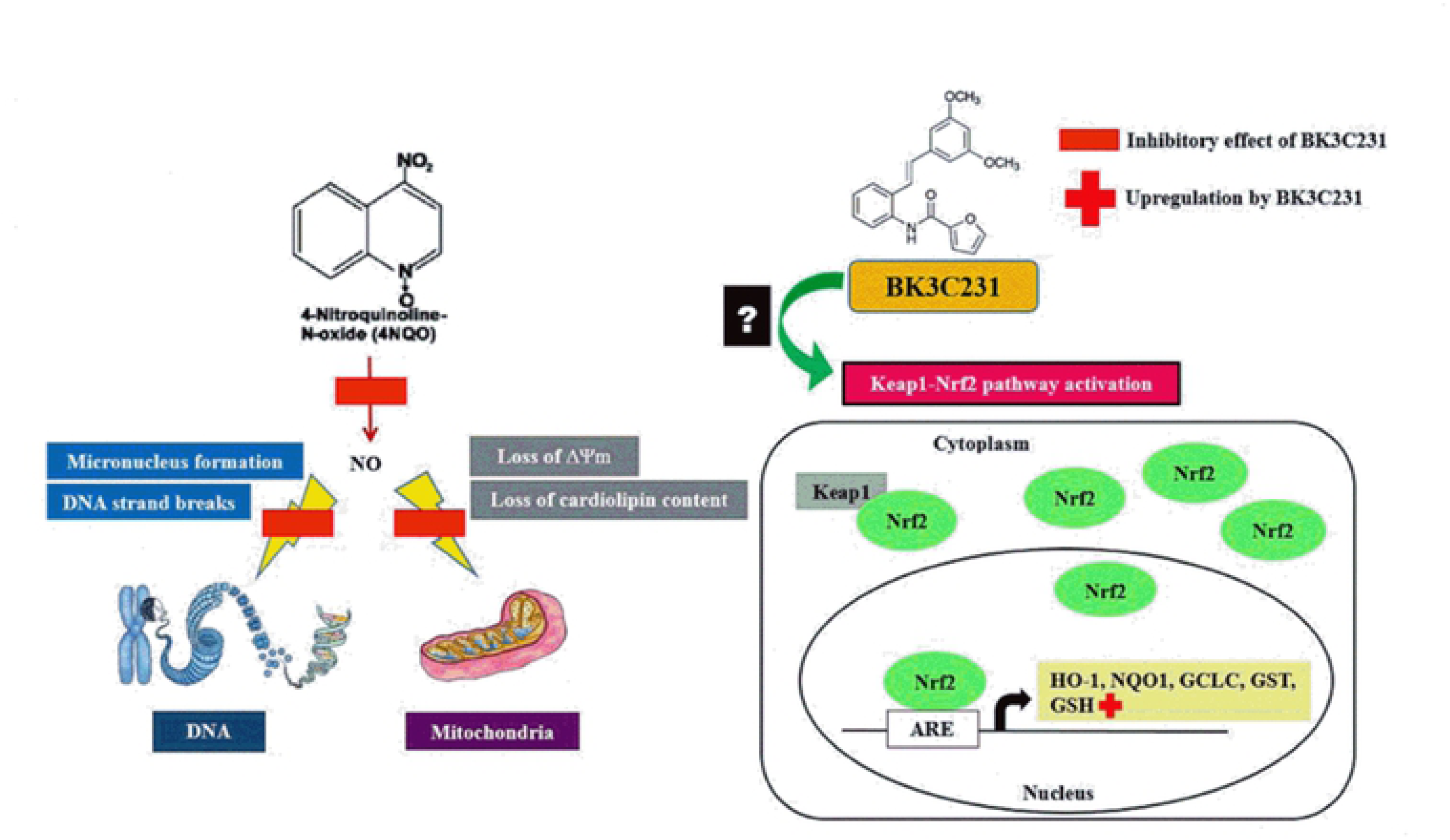
Schematic representation of BK3C231-induced cytoprotection against 4NQO damage in CCD-18Co human colon fibroblast cells. 4NQO caused DNA strand breaks and micronucleus formation as well as mitochondrial membrane potential (ΔΨm) and cardiolipin losses in CCD-18Co cells through NO formation. BK3C231 inhibited these 4NQO-induced DNA and mitochondrial damages by decreasing NO level and increasing GSH level.

## Acknowledgements

The authors would like to thank Dr. Kee Chin Hui from Department of Chemistry, Faculty of Science, University of Malaya for her contribution in the synthesis of compound BK3C231.

## Author contributions

**Conceptualization:** Kok Meng Chan

**Data Curation:** Huan Huan Tan

**Formal Analysis:** Huan Huan Tan

**Funding Acquisition:** Kok Meng Chan

**Investigation:** Huan Huan Tan

**Methodology:** Kok Meng Chan, Huan Huan Tan

**Project Administration:** Kok Meng Chan, Huan Huan Tan

**Resources:** Kok Meng Chan, Noel F. Thomas, Salmaan H. Inayat-Hussain

**Supervision:** Kok Meng Chan, Noel F. Thomas, Salmaan H. Inayat-Hussain

**Validation:** Huan Huan Tan, Kok Meng Chan

**Visualization:** Huan Huan Tan

**Writing – Original Draft Preparation:** Huan Huan Tan

**Writing – Review & Editing:** Huan Huan Tan, Kok Meng Chan

